# Atypical Biological Motion Perception in Children with Attention Deficit Hyperactivity Disorder: Local Motion and Global Configuration Processing

**DOI:** 10.1101/2023.07.25.550597

**Authors:** Junbin Tian, Fang Yang, Ying Wang, Li Wang, Ning Wang, Yi Jiang, Li Yang

## Abstract

Perceiving biological motion (BM) is crucial for human survival and social interaction. Many studies have reported impaired BM perception in autism spectrum disorder, which is characterised by deficits in social interaction. Children with attention deficit hyperactivity disorder (ADHD) often exhibit similar difficulties in social interaction. However, few studies have investigated BM perception in children with ADHD. Here, we compared differences in the ability to process local kinematic and global configurational cues, two fundamental abilities of BM perception, between typically developing and ADHD children. We further investigated the relationship between BM perception and social interaction skills measured using the Social Responsiveness Scale and examined the contributions of latent factors (e.g., sex, age, attention, and intelligence) to BM perception. The results revealed that children with ADHD exhibited atypical BM perception. Local and global BM processing showed distinct features. Local BM processing ability was related to social interaction skills, whereas global BM processing ability significantly improved with age. Critically, general BM perception (i.e., both local and global BM processing) may be affected by sustained attentional ability in children with ADHD. This relationship was primarily mediated by reasoning intelligence. These findings elucidate atypical BM perception in ADHD and the latent factors related to BM perception. Moreover, this study provides new evidence that BM perception is a hallmark of social cognition and advances our understanding of the potential roles of local and global processing in BM perception and social cognitive disorders.

## Introduction

Attention deficit hyperactivity disorder (ADHD) is a common developmental disorder with a prevalence ranging from 2% to 7% in children and adolescents, averaging approximately 5%^1^. In addition to the well-established core symptoms of ADHD (including inability to sustain attention, hyperactivity, and impulsivity), some characteristics of autism spectrum disorder (ASD), such as dysfunction in social communication and social interaction, have also been frequently observed in children with ADHD^2–4^. Nevertheless, experimental studies focusing on social cognition in children with ADHD are limited. Some studies have reported poor performance on social cognition tasks. Among these, impaired theory of mind (ToM) and emotion recognition are the most frequently reported^5–7^. It is difficult for children with ADHD to recognise the emotions and intentions of others. However, our understanding of other social cognitive processes in ADHD remains limited. Further exploration of a diverse range of social cognitions (e.g., biological motion perception) can provide a fresh perspective on the impaired social function observed in ADHD. Moreover, recent studies have indicated that social cognition in ADHD may vary depending on different factors at the cognitive, pathological, or developmental levels, such as general cognitive impairment^5^, symptom severity^8^, or age^5^. Nevertheless, understanding how these factors relate to social cognitive dysfunction in ADHD is still in its infancy. Bridging this gap is crucial as it can help depict the developmental trajectory of social cognition and identify effective interventions for impaired social interaction in individuals with ADHD.

Biological motion (BM), which refers to the movement of a living creature, conveys a wealth of information beyond bodily movements^9^, such as intention^10^, emotion^11^, sex^12^, and identity^13,14^. The advent of point-light display (PLD) technology, which is used to depict human motions,^15^ allows researchers to separate biological motion from other characteristics such as shape and colour. Considering its seminal impact on cognitive, developmental, and clinical neuroscience, BM perception has drawn significant attention from scientists. Some researchers have attempted to deconstruct BM processing into more specific components. Our study concentrated on two fundamental abilities involved in processing BM cues (Figure 1): the ability to process local BM cues derived from major joint motion tracks, and the ability to process global BM cues of human configuration. Previous studies revealed differences between local and global BM perception. Separate neural signals for these abilities imply two independent BM processing stages^16–18^. Local BM cues not only help identify locomotive direction^19^ but also contribute to the detection of life in a visual environment^20^ without the observers’ explicit recognition or attention^21–24^. As a result, the processing of local BM cues is less affected by attention, relatively robust to masking noise, and does not show a learning trend^24,25^. In contrast, global BM processing involves top-down modulation, with attention playing a critical role in its perception^25,26^. Dispersed attention adversely affects performance. Compared with local BM processing, global BM processing is susceptible to learning and is heavily hindered by increased mask densities^25^. These findings suggest that local and global mechanisms play different roles in BM perception, although the exact mechanism underlying this distinction remains unclear. Exploring these two components of BM perception will enhance our understanding of the differences between local and global BM processing and shed light on the psychological processes involved in atypical BM perception.

**Figure 1.**
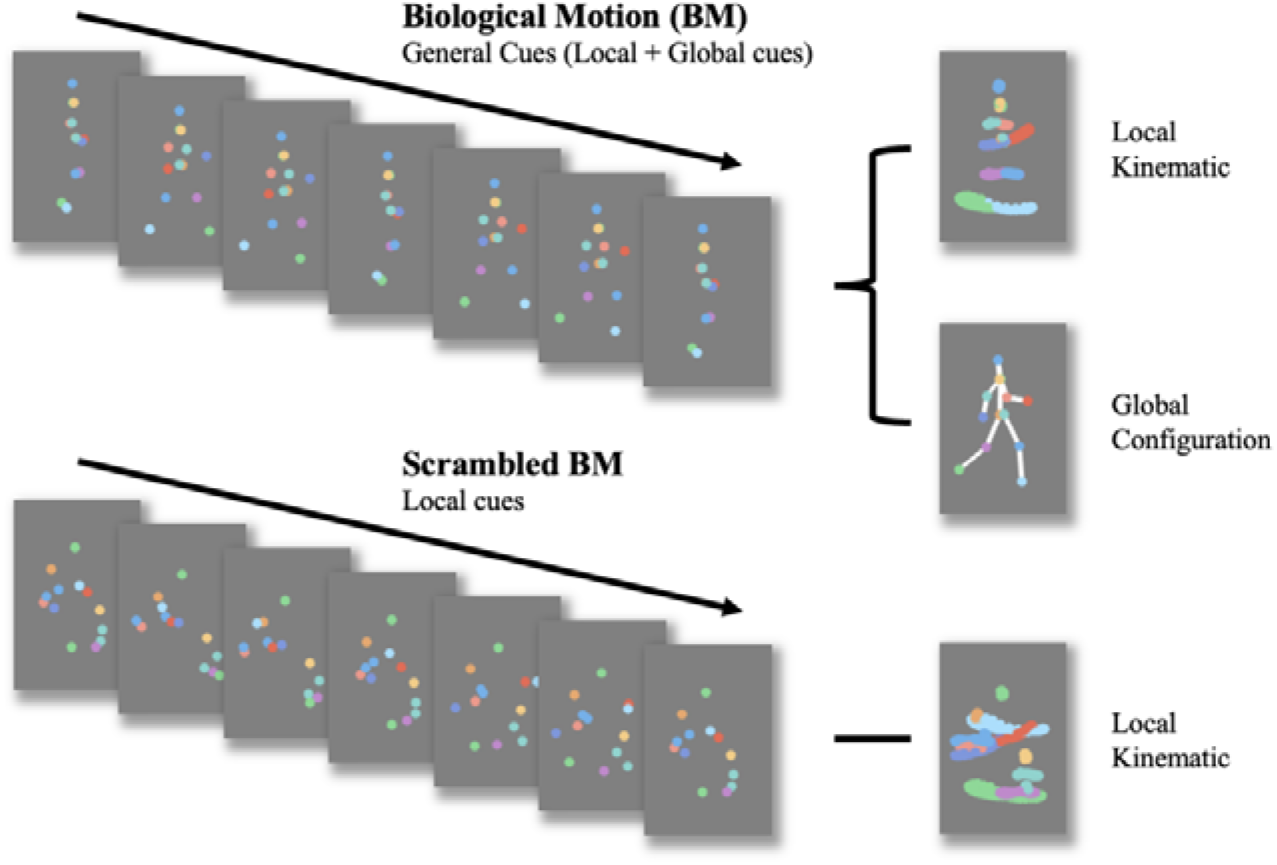
Schematic representation of Biological Motion (BM) and scrambled BM sequence. Intact walker contains information of local kinematics and global configuration. Local kinematics refers to the motion tracks of each critical joint illustrated by the chromatic dot. Global configuration is composed of the relative locations of each joint. In the scrambled BM sequence, global configuration cues have been removed, but local kinematics have been retained. (Figure reconstructed from Wang et al., 2018^30^)

In recent years, BM perception has received significant attention in studies on mental disorders (e.g., schizophrenia^27^) and developmental disabilities, particularly ASD, which is characterised by deficits in social communication and social interaction^28,29^. This is because BM perception is considered a hallmark of social cognition. Individuals with deficits in BM processing exhibit worse social perception in daily life^10^. Another study found that participants’ ability to process BM cues correlated with their autistic traits, particularly in the subdimension of social communication^30^. Therefore, examining BM perception could enhance our understanding of social dysfunction in children with ADHD. Compared with the numerous studies examining impaired BM perception in ASD, few studies have focused on BM perception in children with ADHD. An EEG study found neuroelectrophysiological changes in the processing of BM stimuli in children with ADHD^31^. Specifically, compared with the typically developing (TD) group, children with ADHD showed reduced activity of motion-sensitive components (N200) while watching biological and scrambled motions, although no behavioural differences were observed. Another study found that children with ADHD performed worse in BM detection with moderate noise ratios than the TD group^32^. This finding may be due to the fact that BM stimuli with noise dots will increase the difficulty of identification^33^, which highlights the difference in BM processing between TD and ADHD groups^31^.

Despite initial findings about atypical BM perception in ADHD, previous studies on ADHD treated BM perception as a single entity, which may have led to misleading or inconsistent findings^28^. Hence, it is essential to deconstruct BM processing into multiple components and motion features. To enhance our understanding of the ability to process distinct BM cues in ADHD, we employed a carefully designed behavioural paradigm, as used in our previous study^30^, with slight adjustments made to adapt it for children. This paradigm comprised three tasks: BM-local, BM-global, and BM-general. BM-local assessed the ability to process local BM cues. Scrambled BM sequences were displayed and the participants used local BM cues to judge the direction the scrambled walker was facing. BM-global tested the ability to process the global configuration cues of the BM walker. Local cues were uninformative, and the participants used global BM cues to determine the presence of an intact walker. BM-general tested participants’ ability to process general BM cues (local + global cues). The stimulus sequences consisted of an intact walker and a mask containing similar target local cues so that the participants could use general BM cues to judge the direction the walker was facing.

Experiment 1 examined three specific BM perception abilities in children with ADHD. Children with ADHD show impaired social interaction^2–4^, which implies atypical social cognition. Therefore, we speculated that children with ADHD would perform worse on the three tasks than TD children. In Experiment 2, we further explored the relationship between BM perception and social interaction ability in children with ADHD and identified potential factors (e.g., intelligence quotient [IQ], age, and attention) that may affect BM perception in this population. We speculated that if the mechanisms of processing local and global BM cues are indeed distinct, as suggested by previous studies, then impairment in the ADHD population and the influential factors behind the impairment may be different.

## Materials and methods

### Participants

One hundred seventeen children with and without ADHD were recruited for this study. Eighty-one children met the ADHD diagnostic criteria of the Diagnostic and Statistical Manual of Mental Disorders (DSM-5)^34^. The clinical diagnosis was first made by an experienced child and adolescent psychiatrist in the Child and Adolescent Psychiatric Outpatient Department of Peking University Sixth Hospital, based on the ADHD Rating Scale. The Chinese version of the Kiddie-Schedule for Affective Disorders and Schizophrenia-Present and Lifetime Version DSM-5 (K-SADS-PL-C DSM-5)^35,36^, a semi-structured interview instrument, was then implemented to confirm the diagnosis. Thirty-six TD children from ordinary primary schools in Beijing were screened for the presence of ADHD, ASD, affective disorders, and behavioural disorders by a trained psychiatrist. All participants in the ADHD group had a full-scale IQ >75 (5th upper percentile) on the Wechsler Intelligence Scale for Children-Fourth Edition, and all TD children had a full-scale IQ above the 5th percentile on Raven’s Standard Progressive Matrices^37^, which is used to measure reasoning ability and is regarded as a non-verbal estimate of intelligence. The exclusion criteria for both groups were as follows: (a) neurological diseases; (b) other neurodevelopmental disorders (e.g., ASD, mental retardation, and tic disorders), affective disorders, and schizophrenia; (c) disorders that would impact the completion of the experiment; (d) taking psychotropic drugs or stimulants within the past 30 days; and (e) previous head trauma or neurosurgery.

Thirty-six TD children (age = 9.09 ± 2.18, 14 male) and 39 children with ADHD (age = 9.88 ± 2.23, 28 male) participated in Experiment 1. The groups did not differ by age (*t* = -1.550, *p* = 0.126) but differed in sex (χ*2* = 8.964, *p* = 0.004). Forty-two ADHD children (age = 9.34 ± 1.89, 27 male) participated in Experiment 2. The participants did not participate in Experiment 1. The participants’ demographic characteristics are presented in Table 1. Currently, there is no comparable study on ADHD that indicates effect size as a reference. Studies investigating BM perception in children with ASD typically have sample sizes ranging from 15 to 35 participants per group^29^. Considering the mild impairment of social function in children with ADHD, we determined that a sample size of 35 to 40 participants per group was reasonable for this study. All individuals in each group had normal or corrected-to-normal vision and were naïve to the experimental objectives. Written informed consent was obtained from the parents of all the children before testing. This study was approved by the Institutional Review Boards of Peking University Sixth Hospital and the Institute of Psychology, Chinese Academy of Sciences.

**Table 1.**
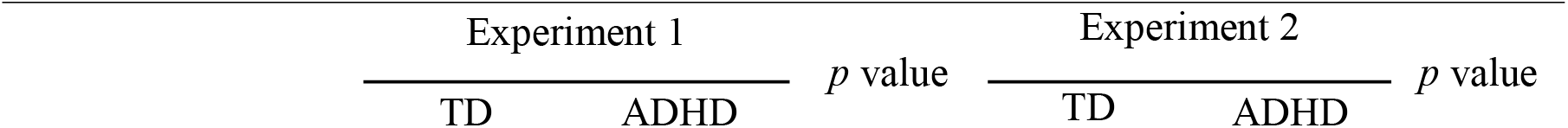
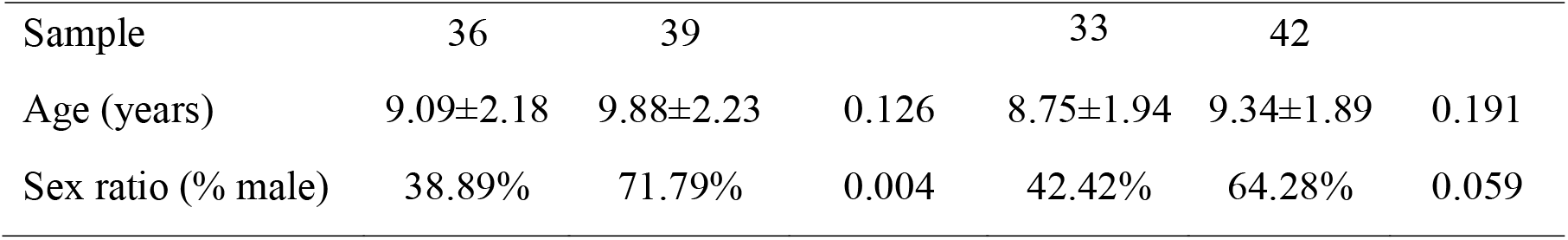
Demographic characteristics of TD and ADHD groups.

### Assessment

#### K-SADS-PL-C DSM-5

The K-SADS-PL-C DSM-5 is a semi-structured interview instrument used to evaluate mental disorders in children and adolescents aged 6–18 years^35^. It involves 35 diagnoses based on the diagnostic criteria of the DSM-5. A trained psychiatrist confirmed the diagnosis by interviewing the parents and children. The Chinese version has demonstrated great psychometric properties^36^.

#### ADHD Rating Scale

The ADHD Rating Scale was adapted from the ADHD diagnostic criteria of the DSM^38^, which requires parents or teachers to complete the scale independently. Its Chinese version has excellent psychometric properties and consists of 2 subscales^39^: inattention (IA, nine items) and hyperactivity-impulsivity (HI, nine items). Each item is rated on a four-point Likert scale ranging from 1 (the symptom appears “never or rarely”) to 4 (the symptom appears “very often”). The final results create three scores: (1) IA dimension score, (2) HI dimension score, and (3) total score. Higher scores indicate more severe ADHD symptoms.

#### Social Responsiveness Scale

The Social Responsiveness Scale (SRS) is a widely used quantitative measure with 65 items used to assess the severity of social impairment in many mental disorders^40^, and the psychometric properties of the Chinese version are reliable^41^. It includes five sub-dimensions: social awareness, social cognition, social communication, social motivation, and autistic mannerisms. Each item is rated on a scale from 0 (never true) to 3 (almost always true), with higher scores indicating worse social ability.

#### Wechsler Intelligence Scale for Children-Fourth Edition

The Wechsler Intelligence Scale for Children-Fourth Edition (WISC-IV) is widely used to test comprehensive intelligence in individuals aged 6–17 years. It contains 15 subtests comprising four broad areas of intellectual functioning: Verbal Comprehension, Perceptual Reasoning, Working Memory, and Processing Speed. The scores in the four broad areas constitute the full-scale intellectual quotient (FIQ).

#### QB Test

The QB Test is a 15-minute Continuous Performance Test (CPT) for assessing inattention and impulsivity, with a high-resolution infrared camera monitoring participant’s activity^42^. Previous psychometric studies have validated its good measurement properties^43^. After the test is completed, several Q scores are calculated to summarise the participants’ performances. The Q scores are standardised based on normative data matched for sex and age. A higher Q score implies more abnormal performance. In this study, we focused on QbInattention, the Q score indicating sustained attention, particularly when children are focused on tasks.

#### Stimuli and Procedure

Point-light BM stimuli sequences adopted in this study have been used in previous studies^44^, which were derived from the configurations of individual walking motions on a treadmill and did not contain an overall translation. Each frame of BM sequences consisted of 13 white dots representing the human head and major joints and was displayed on a grey background (see Video 1). Each walking cycle lasted 1 s with 30 frames. For each trial, the initial frame of BM sequences was randomised. The entire point-lighted BM walker was presented at approximately a 5.7° vertical visual angle. Stimuli were presented on a 14-inch monitor, and responses were evaluated using MATLAB together with PsychToolbox extensions. All subjects completed the experiments in a dimly lit room with their heads on a chinrest to ensure that their eyes were 50 cm away from the monitor.

In Experiment 1, the children were required to complete three tasks that were similar to but slightly modified from the versions implemented in our previous study^30^ (Figure 2). Each trial began with a fixation cross (0.6° * 0.6°). Following a random interval of 1200-1500 ms, the monitor displayed a task-specific BM sequence lasting for 2 s (60 frames).

**Figure 2.**
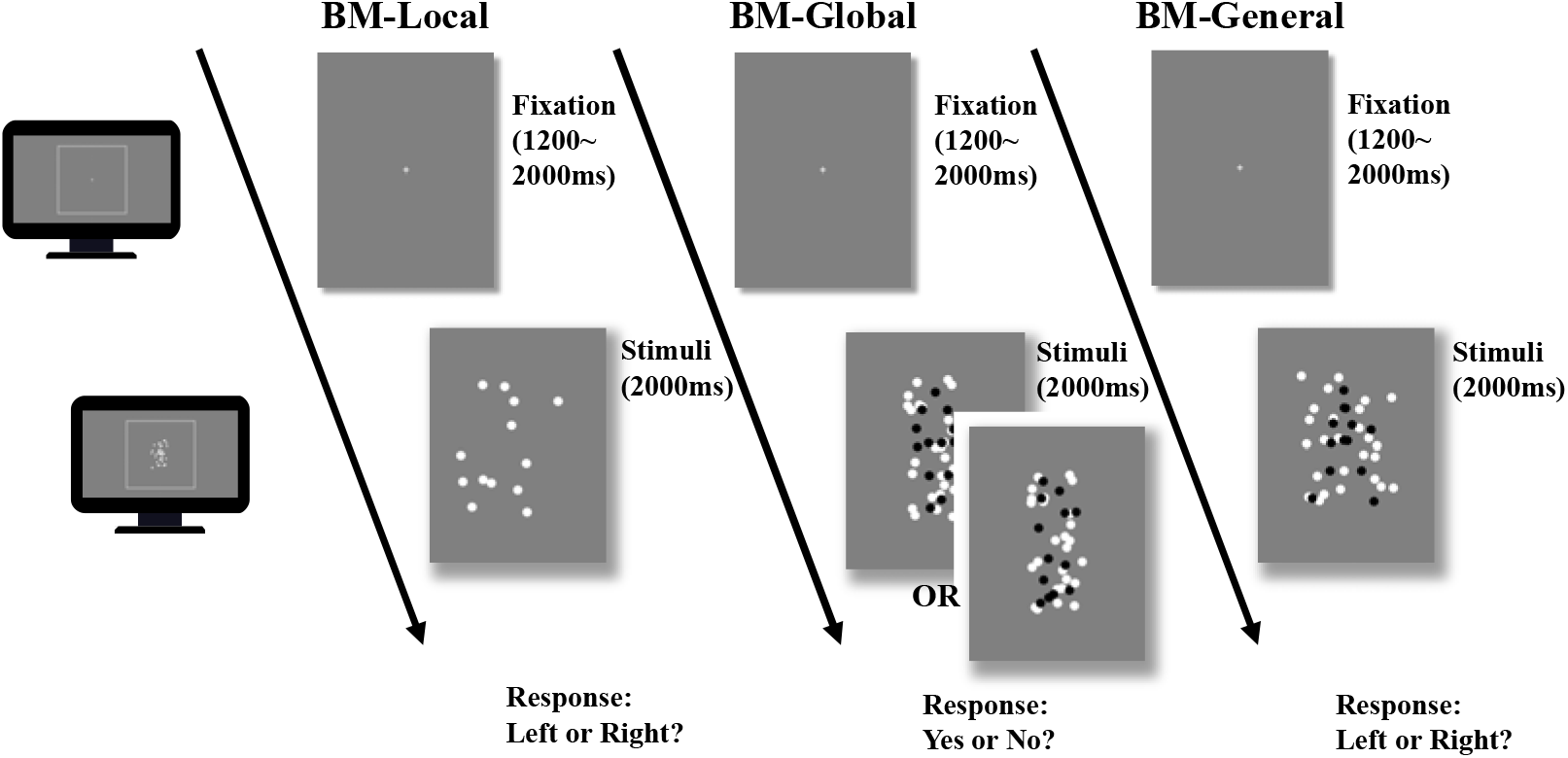
Illustration of the trial sequence. In BM-local, a monitor displayed scrambled BM sequences. Participants only judged the facing direction of the scrambled walker using local BM cues. In BM-global, each trial only showed an intact or scrambled walker (black dots in the figure) embedded within a mask containing local BM cues. Because the two conditions contained the same local cues that were also present in the mask, the participant must rely on global BM cues to determine whether an intact walker was present in the mask. The figure shows one of five possible directions the intact walker could face (i.e., facing participants). In BM-general, the stimuli sequence consisted of an intact walker (black dots) and a mask containing similar target local cues, and children judged the direction the walker was facing using general BM cues (local + global). Dots in the figure are rendered in black for better illustration but were displayed in white in the actual experiments.

BM-Local assessed participants’ ability to process local BM cues. During the task, the monitor displayed only a scrambled walker facing either the left or right (Video 2). Specifically, the 13 dots constituting the intact walker were randomly relocated within the original range of the BM walker (randomly presented in 2D). This manipulation disrupted the global configuration of the intact walker while retaining local kinematics. After the display, we required the children to press a left or right button to indicate the direction of motion of the unidentified creature (i.e., the scrambled BM walker) as accurately as possible. Children did not receive feedback on the accuracy of each response. Thirty trials were conducted, with 15 trials for each condition (left and right).

BM-Global tested the ability to process the global configuration cues of the BM walker. A target walker (scrambled or intact) was displayed within the mask (Video 3) during this task. The mask consisted of two scrambled target walkers (26 dots) with the same locomotion direction as the target walker, displayed within a boundary approximately 1.44 times larger than the intact walker. A scrambled or intact version of the target walker was randomly embedded in the mask and entirely overlaid. Thus, the global BM component could be isolated as two conditions (i.e., scrambled and intact walkers) containing the same local kinematics information, rendering the local motion cues uninformative. The children were required to judge whether there was an intact walker on the mask. A correct response relied on the extraction of global cues from an intact walker. To prevent children from learning the shape of the walker^21^, we set target walkers that possibly faced one of five equally spaced directions from left to right. Of the five walkers used, two faced straight to the left or right, orthogonal to the viewing direction. Two walked with their bodies oriented at a 45-degree angle to the left or right of the observer. The last one walked towards the observer. Video 4 shows the five facing directions of the walker. Thirty trials were conducted consisting of two conditions (intact or scrambled target).

BM-General tested participants’ ability to process general BM cues (local and global). In BM-General, the monitor displayed an intact walker (facing either the left or right) embedded within a mask (see Video 5). The mask used in this task was similar to that used in BM-Global. The children were required to judge the direction the target walker was facing (left or right). Because the mask and target walker contained the same local BM cues and the target walker was presented with additional global configuration cues, children could rely on general BM information (i.e., a combination of local and global cues) to perform the task. BM-General consisted of 30 trials, with 15 trials for each facing direction. The other parameters of BM-Global and BM-General were similar to those of BM-Local. Before each task, the children practiced for five trials to ensure a good understanding. We performed the three tasks in a fixed order so that the participants were naïve to the nature of the local BM cues in BM-Local.

In Experiment 2, 42 children with ADHD completed the same procedure as in Experiment 1. In addition, the parents completed the SRS to assess social interaction.

### Statistics

Two-sample t-tests were used to examine the difference in BM perception abilities between TD and ADHD children^45,46^, and Pearson’s correlation analyses were used to assess the relationship between the accuracy of each task and the SRS score.

Additionally, general linear models and path analyses were used to explore potential factors influencing BM perception. A *p*-value < 0.05 was considered statistically significant. Path analyses were conducted using AMOS, whereas other analyses were conducted using SPSS.

## Results

### Children with ADHD exhibit atypical BM perception

Figure 3 displays the mean accuracies (ACC) for both the TD and ADHD groups across the three tasks in Experiment 1. We examined the difference in the ACC between the TD and ADHD groups for each task using a two-sample t-test. The results of BM-Local showed a significant difference (TD: 0.52 ± 0.13, ADHD: 0.44 ± 0.09, *t*_73_ = 3.059, *p* = 0.003, Cohen’s *d* = 0.716), indicating that children with ADHD exhibited impaired local BM processing ability. For BM-Global and BM-General, where children were asked to detect the presence or discriminate the direction the target walker was facing, the TD group had higher accuracies than the ADHD group (BM-Global - TD: 0.70 ± 0.12, ADHD: 0.59 ± 0.12, *t*_73_ = 3.677, *p* < 0.001, Cohen’s *d* = 0.861; BM-General - TD: 0.79 ± 0.12, ADHD: 0.63 ± 0.17, *t*_73_ = 4.702, *p* < 0.001, Cohen’s *d* = 1.100). These findings suggest the presence of impaired global and general BM perception in children with ADHD. To ensure that sex did not influence the results, we conducted a subsampling analysis with balanced data, and the results remained consistent (see Supplementary Information).

**Figure 3.**
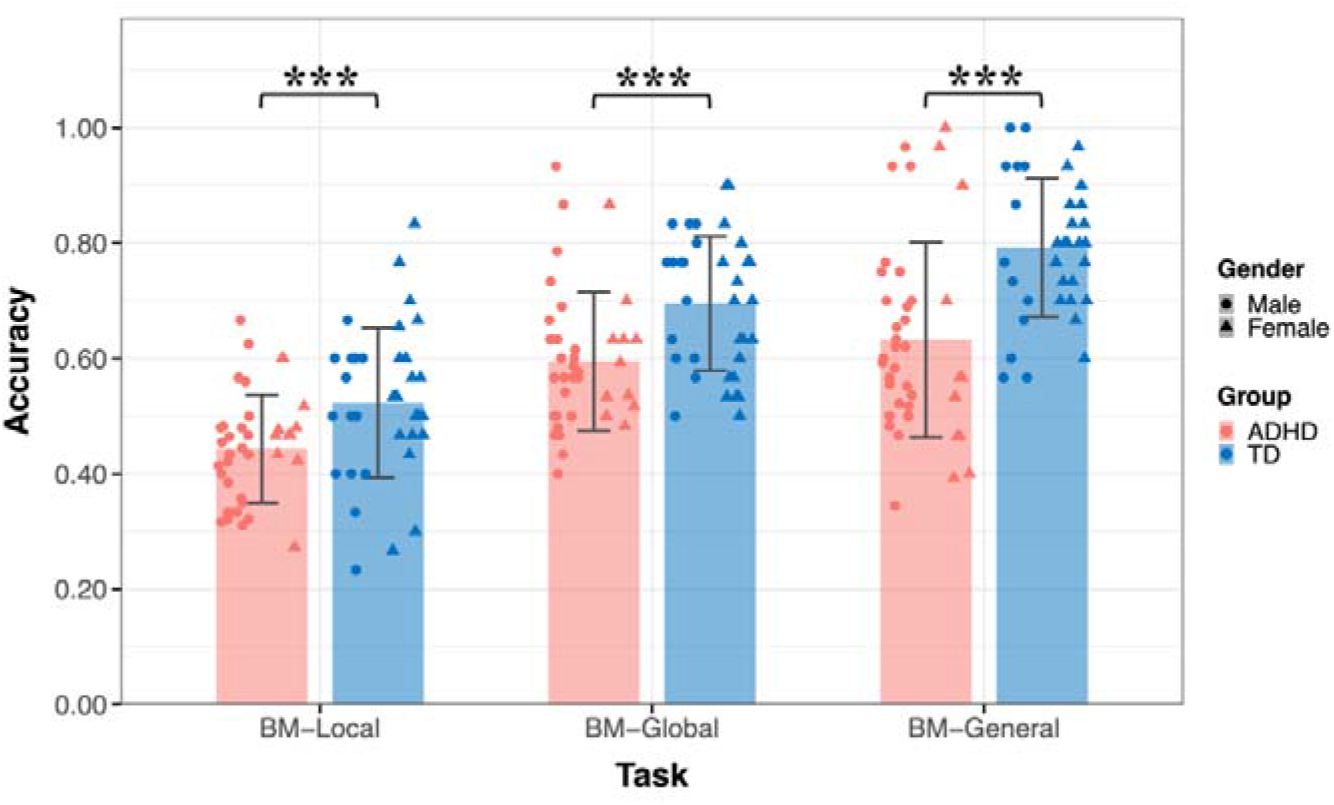
The mean accuracy of the three tasks. Typically developing (TD) children had higher accuracies than children with attention deficit hyperactivity disorder (ADHD) in the three tasks in Experiment 1. Error bars show standard deviations. \*\*\**p* < 0.001

### Atypical perception of local BM information predicts impaired social interaction in ADHD

Experiment 1 provides evidence of atypical BM perception in children with ADHD. Previous studies have revealed that BM processing ability is a hallmark of social cognition^10^ and is negatively correlated with social ability^30^. Substantial evidence indicates that children with ADHD often experience problems with social interaction. We hypothesised that compromised social interaction in children with ADHD would be associated with BM processing. To confirm this hypothesis, we recruited 42 naïve children with ADHD to participate in Experiment 2 and examined the relationship between their social interaction abilities and BM perception. Parents or caregivers completed the SRS. The SRS total score of the ADHD group was higher than that of the TD group (SRS total score - ADHD: 54.64 ± 18.42, TD: 38.64 ± 12.47, *t* = -4.277, *p* < 0.001). We found that children with higher total SRS scores performed worse on the three tasks; that is, the abilities of BM processing were negatively correlated with SRS total score (BM-Local: *r* = -0.264, false discovery rate [FDR]-corrected *p* = 0.033; BM-Global: *r* = -0.238, FDR-corrected *p* = 0.039; BM-General: *r* = -0.359, FDR-corrected *p* = 0.006).

The correlations encompassing all data from both groups might reflect group disparities, given the significant distinction in SRS total score between TD and ADHD children, alongside their marked differences in BM processing abilities. Therefore, we conducted additional subgroup analysis to further explore the relationship between social interaction and BM processing ability in children with ADHD. As depicted in Figure 4, the correlation between the SRS total score and the ability to process local cues was only found in the ADHD group (ADHD: *r* = -0.461, FDR-corrected *p* = 0.004; TD: *r* = 0.109, FDR-corrected *p* = 0.547), particularly on subscales of social awareness, social cognition, social communication, social motivation (see Table 2 for detailed information). However, we did not find a statistically significant correlation between the SRS total score and global or general BM processing in either the ADHD or TD groups (global BM perception, TD: *r* = - 0.020, FDR-corrected *p* = 0.910, ADHD: *r* = -0.207, FDR-corrected *p* = 0.374; general BM perception, TD: *r* = -0.118, FDR-corrected *p* = 0.514, ADHD: *r* = -0.286, FDR-corrected *p* = 0.134).

**Figure 4.**
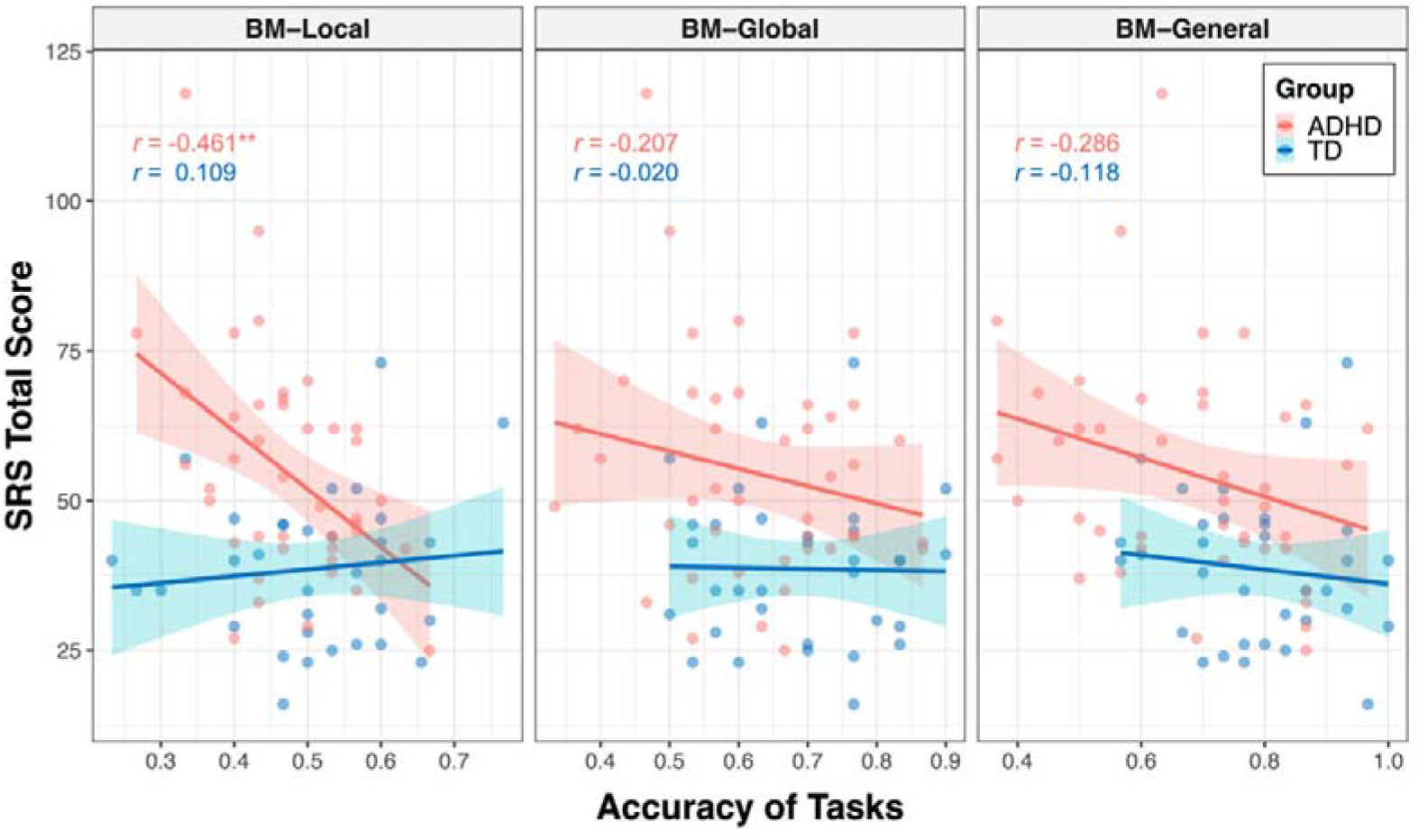
Correlations between response accuracies and SRS total score. The ability to process local cues is significantly correlated with the SRS total score in the ADHD group. The grey shading represents the 95% confidence interval. ** FDR-corrected *p* < 0.01.

**Table 2.** The correlation between the ability of local BM processing and subdimensions of SRS in ADHD children.

|  | Correlation coefficient (r) | FDR-corrected *p* value |
| --- | --- | --- |
| Social awareness | -0.333 | 0.039 |
| Social cognition, | -0.416 | 0.020 |
| Social communication | -0.381 | 0.022 |
| Social motivation | -0.406 | 0.020 |
| Autistic mannerisms | -0.245 | 0.117 |

To determine the specificity of the correlation between local BM processing and SRS total score in the ADHD group, we constructed general linear models to further compare these correlations (see Supplementary Information). We observed a significantly stronger correlation between SRS total score and response accuracy in the ADHD group compare to the TD group (*p* = 0.003) for the BM-Local, but not for the BM-Global (*p* = 0.381) or BM-General (*p* = 0.455). Additionally, our results showed trends towards significance, indicating that the correlation between the SRS total score and response accuracy of the ADHD group in BM-Local was more negative than that in BM-Global (*p* = 0.074) or in BM-General (*p* = 0.073). These findings suggest that the atypical local BM processing ability may be specifically related to the impairment of social interaction in children with ADHD.

### Global BM processing develops with age and is regulated by reasoning intelligence and attention function

Many factors can affect social cognition in ADHD, such as general cognitive impairment^5^, symptom severity^8^, and age^5^. To better understand their role in atypical BM processing, we examined the relationship between BM task performance and factors such as age, IQ, and attention function in the ADHD group. Data from the ADHD group in both Experiments 1 and 2 were integrated, resulting in 80 ADHD participants (one child did not complete the QB test). Three linear models were built to investigate the contributing factors: (a) *ACC_BM-Local_ =* β*_0_ +* β*_1_ * age +* β*_2_ * gender +* β*_3_ * FIQ +* β*_4_* QbInattention*, (b) *ACC_BM-Global_ =* β*_0_ +* β*_1_ * age +* β*_2_ * gender +* β*_3_ * FIQ +* β*_4_ * QbInattention*, and (c) *ACC_BM-General_ =* β*_0_ +* β*_1_ * age +* β*_2_ * gender +* β*_3_ * FIQ +* β*_4_ * QbInattention +* β*_5_* ACC_BM-Local_ +* β*_6_ * ACC_BM-Global_*. *ACC_BM-Local_*, *ACC_BM-Global_* and *ACC_BM-General_* refer to the response accuracies of the three tasks in the ADHD group, and *QbInattention* is the standardised score for sustained attention function. We screened the factors with the largest contribution to the models using stepwise regression. In model (a), no variable remained after stepwise regression, suggesting that local BM processing remained stable with age and was not affected by attention or IQ. In model (b), the ability to process global BM cues was enhanced with age (standardised β*_1_* = 0.251, *p* = 0.025). In model (c), higher FIQ, particularly on the subdimension of Perceptual Reasoning (standardised β*_3_* = 0.271, *p* = 0.005), and better performance in global BM processing (standardised β*_6_* = 0.290, *p* = 0.004) predicted better performance in general BM processing. Furthermore, as children aged, the ability to probe general BM information improved (standardised β*_1_* = 0.365, *p* < 0.001). It is worth noting that QbInattention showed a strong negative correlation with Perceptual Reasoning (*r* = -0.355, *p* = 0.001) and general BM perception (*r* = - 0.246, *p* = 0.028). Owing to the potential collinearity issue, we employed a post hoc path analysis to visualise these relationships (Figure 5). The results indicated that sustained attention (i.e., QbInattention) did not directly predict performance in BM-General but was significantly indirectly predicted by Perceptual Reasoning ability.

**Figure 5.**
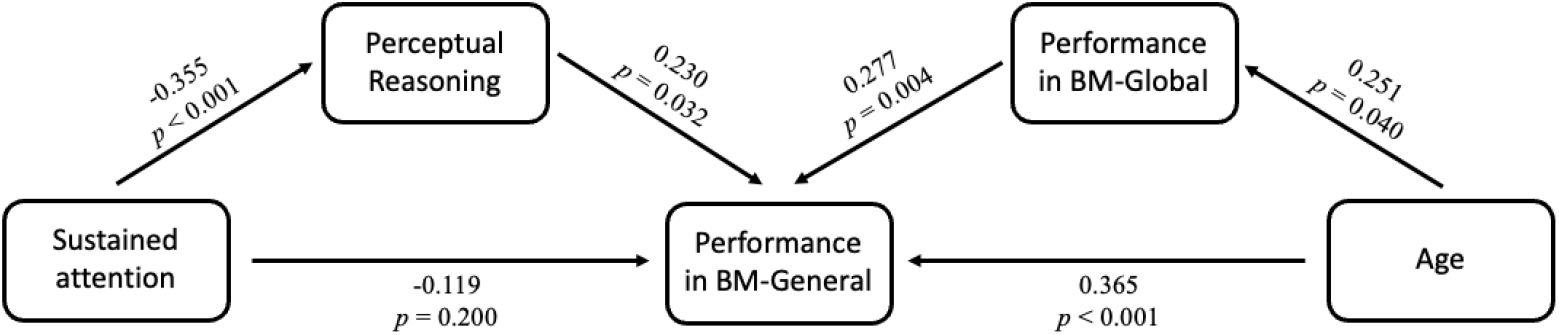
Factors influencing BM perception in ADHD children. Post hoc path analysis confirmed that the effect of sustained attention on performance in BM-General was entirely mediated by Perceptual Reasoning, and the ability of global BM processing partly mediated the effect of age on performance in BM-General.

Furthermore, as children with ADHD aged, their performance in BM-General improved, both directly and through the enhanced processing of global BM cues. We also built three models to explore further the effects of Reasoning IQ and age on BM perception in TD children: (d) *ACC_BM-Local_ =* β*_0_ +* β*_1_ * age +* β*_2_ * gender +* β*_3_ * FIQ*. (e) *ACC_BM-Global_ =* β*_0_ +* β*_1_ * age +* β*_2_ * gender +* β*_3_ * FIQ*; (f) *ACC_BM-General_ =* β*_0_ +* β*_1_ * age +* β*_2_ * gender +* β*_3_ * FIQ +* β*_4_ * ACC_BM-Local_ +* β*_5_ * ACC_BM-Global_*. In model (d), no regressor remained significant after stepwise regression. However, in models (e) and (f), we observed positive relationships between age and performance (model e: standardized β*_1_* = 0.396, *p* = 0.017; model f: standardised β*_1_* = 0.330, *p* = 0.049). We also conducted a path analysis similar to that in the ADHD group and found no statistically significant mediator effect (Figure S3). The complete information about models a-f could be found in Table S1.

In summary, our findings suggest that the ability to perceive global and general BM cues, rather than local BM cues, improves with age in both groups. We speculated that age-related improvement was different when processing different BM cues and in different groups. Therefore, we examined the differences between the age-related improvement in processing local cues and that in processing global and general cues (see Supplementary Information for detailed methods). In the ADHD group, we observed that the ability to process general BM cues significantly improved with age compared to local cues (*p* < 0.001) and a trend that the improvement in processing global BM cues with age was greater than that in processing local BM cues (*p* = 0. 073). However, these patterns were not observed in the TD group (see Supplementary Information). In addition, we examined the differences in the improvements in processing of BM cues with age between the two groups for each task. There was no difference in the effect of age on the response accuracy between the TD and ADHD groups for the three tasks (see Supplementary Information).

## Discussion

Our study contributes several promising findings concerning atypical BM perception in children with ADHD. Specifically, we observed atypical local and global BM perception in children with ADHD. Notably, local and global BM processing exhibited distinct features. The ability to process local BM cues appears to be associated with social interaction traits in children with ADHD. In contrast, global BM processing was associated with age-related development. In addition, general BM perception may be affected by factors such as attention.

BM perception is a widely studied topic in visual cognition owing to its inherent biological and social properties. BM processing has significant value in successfully navigating daily life, particularly in non-verbal communication^11,47^ and adaptive behavior^26,48^. In TD children, there is a clear association between BM perception and social cognitive abilities^49^. For example, 12-month-old infants exhibit social behaviours (i.e., following gaze) elicited by BM displays^50^. Therefore, BM perception plays a crucial role in the development of children’s social cognition. This ‘social interpretation’ of BM suggests that the difficulties in processing BM may serve as an indicator of impaired social interaction^30^. Our results are consistent with these findings. We observed atypical BM perception in children with ADHD and a significant relationship between BM perception performance and the SRS total score. Further subgroup analysis revealed a significant negative correlation between SRS total score and the accuracy of local BM processing in the ADHD group. This correlation was stronger in the ADHD group than in the TD group. The lack of a significant correlation may be due to the narrow range of SRS scores in the TD group. Future studies should increase the sample size to explore the correlations among diverse individuals. These findings suggest that BM processing is a distinct hallmark of social cognition in ADHD children^10,30^.

BM perception is a multi-level phenomenon^51–53^. At least in part, the processing of local and global BM information appears to involve different neural mechanisms^16^. Sensitivity to local BM cues emerges early in life^54,55^ and involves rapid processing in the subcortical regions^16,56–58^. As a basic pre-attentive feature^23^, local BM cues can guide visual attention spontaneously^59,60^. In contrast, the ability to process global BM cues is related to slow cortical BM processing and is influenced by many factors such as attention^25,26^ and visual experience^21,51^. As mentioned above, we found a significant negative correlation between the SRS total score and the accuracy of local BM processing, specifically in the ADHD group. This could be due to decreased visual input related to atypical local BM processing, which further impairs global BM processing. According to the two-process theory of biological motion processing^61^, local BM cues guide visual attention toward BM stimuli^55,62^. Consequently, the visual input of BM stimuli increases, facilitating the development of the ability to process global BM cues through learning^21,63^. The latter is a prerequisite for attributing intentions to others and facilitating social interactions with other individuals^20,64,65^. Thus, atypical local BM processing may contribute to impaired social interactions through altered visual input. Further empirical studies are required to confirm these hypotheses.

The ability to process global BM cues develops with age. Previous studies indicated that global BM perception is enhanced with age in TD children^66,67^. This developmental trend in global BM processing is also evident in individuals with impaired BM perception (e.g., children with ASD). BM processing performance in children with ASD becomes more aligned with that of TD children as they age^29,68–70^. Our study contributes new evidence to the understanding of the development of global BM processing. We found that the ability to process global and general BM cues improved significantly with age in both the TD and ADHD groups, implying that the processing module for global BM cues tends to mature with development. This finding is akin to the potential age-related improvements observed in certain aspects of social cognitive deficits in individuals with ADHD^5^. Interestingly, in the ADHD group, the improvement in processing general and global BM cues was greater than in processing local BM cues. Few developmental studies have been conducted on local BM processing. The ability to process local BM cues remained stable and did not exhibit a learning trend^21,25^. A reasonable interpretation may be that local BM processing is a low-level mechanism, probably performed by the primary visual cortex and subcortical regions such as the superior colliculus, pulvinar, and ventral lateral nucleus^14,56,61^, which are not malleable. Although there was no statistical difference between the improvements in processing local and global BM cues in the TD group, this may be due to the relatively small sample size or the relatively higher baseline abilities of BM perception in TD children, resulting in a relatively milder improvement. It is worth noting that the ability to process global BM cues was positively correlated with the performance in processing general BM cues in the ADHD group, whereas no such correlation was found in the TD group. This suggests that TD children are able to extract and integrate both local and global cues, whereas children with ADHD may rely more on global BM cues to judge the direction the walker is facing when presented with both local and global BM cues, which correspond to a hierarchical model^51^. Once a living creature is detected, an agent (i.e., is it a human?) can be recognised by a coherent, articulated body structure that is perceptually organised based on its motions (i.e., local BM cues)^71^. This involves top-down processing and probably requires attention^25,72^, particularly in the presence of competing information^26^. Our findings are consistent with those of previous studies on the cortical processing of BM^73^, as we found that the severity of inattention in children with ADHD was negatively correlated with their performance in global BM processing, whereas this significant correlation was not found in local BM processing, which may involve bottom-up processing^61,65^ and might not need participants’ explicit attention^21,23,74,75^. However, further studies are needed to verify this hypothesis.

Interestingly, children with impaired BM perception may employ a compensatory strategy^76^. Previous studies found no impairment in BM recognition in individuals with autism with high IQ^76^, but children with ASD exhibited weaker adaptation effects for biological motion than TD children^77^. One possibility is that individuals with a high IQ and impaired BM perception can develop or employ reliable strategies for BM recognition, compensating for the lack of intuitive social perceptual processing^78,79^. The current study supports this assumption, as children with a higher IQ, particularly in Perceptual Reasoning, demonstrated better performance. Owing to the impact of attention deficits on Perceptual Reasoning, the performance of children with ADHD did not align with that of TD children.

Overall, our study reveals two distinct and atypical fundamental abilities underlying BM perception in children with ADHD. Notably, anomalous local BM processing may predict impaired social interactions in children with ADHD. Moreover, these results revealed the potential contributions of age, IQ, and attention to BM information processing. These findings also shed new light for future studies. First, different developmental trends appear in local and global BM processing. Further studies are required to explore the relationship between the two fundamental BM processing abilities, which will contribute to understanding the respective and mutual neural mechanisms underlying the two types of BM processing. Second, exploring the performance of more advanced BM processing in children with ADHD, such as emotion and identity recognition in BM tasks, is necessary to delineate the neural profiles involved in processing BM in ADHD. Finally, a comparative study between ADHD and ASD is warranted to identify common neuropsychological traits and biomarkers of social cognition impairment.

## Acknowledgements

We thank the three reviewers for their constructive comments, and Dr. Shuo Gao and Dr. Li Shen for assistance with manuscript revision. Special thanks to the parents and the children who took part in the study. This research was supported by grants from the Beijing Municipal Science and Technology Commission (Z181100001518005), the Ministry of Science and Technology of China (2021ZD0203800), the National Natural Science Foundation of China (31830037), the Interdisciplinary Innovation Team (JCTD-2021-06), and Fundamental Research Funds for the Central Universities.

## Author contributions

Junbin Tian: Conceptualization, Data curation, Formal analysis, Investigation, Methodology, Visualization, Project administration, Writing – original draft. Fang Yang: Formal analysis, Methodology, Investigation. Ying Wang: Methodology, Software, Writing – review & editing. Li Wang: Methodology, Software, Writing – review & editing. Ning Wang: Investigation. Yi Jiang: Conceptualization, Methodology, Supervision, Writing – review & editing. Li Yang: Conceptualization, Methodology, Supervision, Writing – review & editing.

## Declaration of interests

The authors declare no competing interest.

## Ethics

All procedures contributing to this work comply with the ethical standards of the relevant national and institutional committees on human experimentation and with the Helsinki Declaration of 1975, as revised in 2008. The institutional review boards of the Peking University Sixth Hospital and the Institute of Psychology, Chinese

Academy of Sciences have approved this study (reference number for approval: (2020) Ethics Review No.9 and H23030).

## Data availability

The data analyzed during the study is available at https://osf.io/37p5s/

## Notes

### Competing Interest Statement

The authors have declared no competing interest.

### Summary of Updates

New version of manuscript has been revised according to the suggestion from eLife's assessment.

