## Supplementary Information for "Atypical Biological Motion Perception in Children with Attention Deficit Hyperactivity Disorder: Local Motion and Global Configuration Processing"

**Children with ADHD exhibit** **atypical BM** **perception** **(subsampled balanced data)**

To address potential gender-related confounds, we applied a statistical matching technique, Propensity Score Matching (PSM)^1^, to obtain a sub-dataset (1:1 matching) using the Matching package in R 4.3.0. We matched the characteristics, namely age and gender, to create more similar groups. After matching, 24 TD children (age = 9.70 ± 2.08, 14 male) and 24 children with ADHD (age = 9.67 ± 1.93, 14 male) were included in the following analysis. The two groups did not differ significantly in age (*t* = 0.041, *p* = 0.967) and had an identical gender ratio. Our findings, as reproduced in Figure S1, confirmed that the TD group exhibited higher response accuracy than the ADHD group in all three tasks (BM-Local - TD: 0.51 ± 0.11, ADHD: 0.45 ± 0.10, *t*_46_ = 2.092, *p* = 0.042, Cohen's *d* = 0.617; BM-Global - TD: 0.73 ± 0.11, ADHD: 0.57 ± 0.10, *t*_46_ = 5.116, *p <* 0.001, Cohen's *d* = 1.51; BM-General - TD: 0.79 ± 0.14, ADHD: 0.58 ± 0.15, *t*_46_ = 5.088, *p <* 0.001, Cohen's *d* = 1.50).

**The correlation between SRS total scores and the ability to process local BM cues in the ADHD group is significantly stronger than in the TD group**

To further determine the specificity of the correlation between local BM processing and SRS total score in the ADHD group. We constructed general linear models to compare these correlations. We first examined the difference in the correlations between the response accuracy for each task and SRS total score between the TD and ADHD groups. The differences between correlations were tested by testing interaction terms in general linear models. Specifically, we recoded the categorical variable (i.e., *group*) into a dummy variable (*D*) using the TD group as a reference and constructed a general linear model for each task (Table S2, Model 1-3): *SRS =* *β_0_ + β_1_ * ACC + β_2_ * d + β_3_ * (ACC * D)*. *ACC* refers to the response accuracy, and *SRS* refers to the SRS total score. If the effect of the interaction term (i.e., *β_3_*) is statistically significant, it indicates that the correlations between response accuracy and SRS total score for the TD and ADHD groups are significantly different^2^. For BM-Local, we observed that the correlation between SRS total score and ACC in the ADHD group was significantly stronger than in the TD group (standardized *β_3_* = -0.629, *p* = 0.003). However, no significant difference was identified with regard to BM-Global (standardized *β_3_* = -0.195 *p* = 0.381) or BM-General (standardized *β_3_* = -0.179, *p* = 0.455).

In addition, we further examined the relative differences in correlations with SRS total score between BM-Local and BM-Global or BM-General in the ADHD group (Figure S2). Similarly, we recoded three task types to two dummy variables, *D_1_* and *D_2_*_,_ using BM-Local as a reference. The coefficient of *D_1_* represents the difference in relationship to SRS total score between BM-Local and BM-Global, and the coefficient of *D_2_* represents the difference in relationship to SRS total score between BM-Local and BM-General. A general linear model was constructed (Table S2, Model 4): *SRS = β_0_ + β_1_ * ACC + β_2_ * D_1_ + β_3_ * D_2_ + β_4_ * (ACC * D_1_) + β_5_ * (ACC * D_2_)*. If the effect of the interaction term (i.e., *β_4_* or *β_5_* ) is statistically significant, it indicates a difference in correlations with SRS total score between BM-Local and BM-Global (or BM-General). The results suggested trends where the correlations with SRS total score were more negative for BM-Local relative to BM-Global (standardized *β_4_* = 0.580 p = 0.074) and BM-General (standardized *β_5_* = 0.550 p = 0.073).

**The improvement in processing general BM cues with age is significantly greater than in processing local BM cues in ADHD group**

To further examined the differences between the age-related improvement in processing local cues and that in processing global and general cues (Figure S4), we employed similar analyses as described earlier. We recoded task types into two dummy variables, *D_1_* and *D_2_*, using BM-Local as a reference. The coefficient of *D_1_* represents the difference in relationship to age between BM-Local and BM-Global, and the coefficient of *D_2_* represents the difference in relationship to age between BM-Local and BM-General. The following model was created for each group (Table S3, Model 5-6): *ACC = β_0_ + β_1_ * age + β_2_ * D_1_ + β_3_ * D_2_ + β_4_ * (age * D_1_) + β_5_ * (age * D_2_)*. If the effect of the interaction term (i.e., *β_4_* or *β_5_*) is statistically significant, it indicates a difference in the effect of age on ACC between BM-Local and BM-Global (or BM-General). In the ADHD group, we observed a significant difference in the effect of age on ACC between BM-Local and BM-General (standardized *β_5_* = 0.462, *p* < 0.001) and marginally significant differences in the effect of age on ACC between BM-Local and BM-Global (standardized *β_4_* = 0.228, *p* = 0.073). However, there was no difference in the effect of age between BM-Local and BM-Global or BM-General in the TD group (standardized *β_4_* = 0.095, *p* = 0. 575; standardized *β_5_* = 0.056, *p* = 0. 739).

We also examined the differences in age-related improvements in processing BM cues between two groups for each task (Figure S5). We recoded the variable *group* into a dummy variable (*D*) using the TD group as a reference, and established a general linear model for each task (Table S3, Model 7-9): *ACC = β_0_ + β_1_ * age + β_2_ * D + β_3_ * (age * D)*. If the effect of the interaction term (i.e., *β_3_*) is statistically significant, it indicates a difference in the effect of age on ACC between the TD group and the ADHD group. The results showed no difference in the effect of age on ACC between the TD and the ADHD group in three tasks (BM-Local: standardized *β_3_* = -0.306, *p* = 0.112; BM-Global: standardized *β_3_* = -0.091, *p* = 0.621; BM-General: standardized *β_3_* = 0.192, *p* = 0.263).

**Figures**

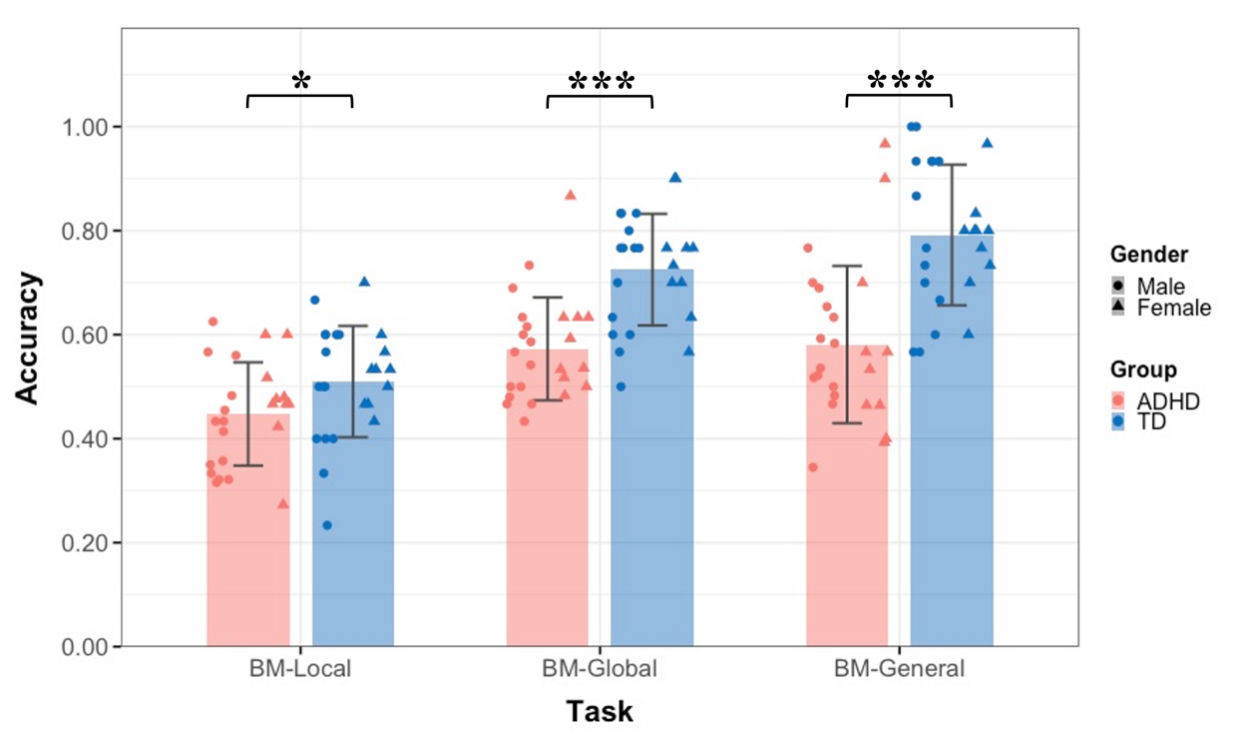

**Figure S1. The mean accuracy of the three tasks (subsampled balanced data).** Typically developing (TD) children had higher accuracies than children with attention deficit hyperactivity disorder (ADHD) in three tasks in Experiment 1. Error bars show standard deviations. **p* < 0.05; ****p* < 0.001

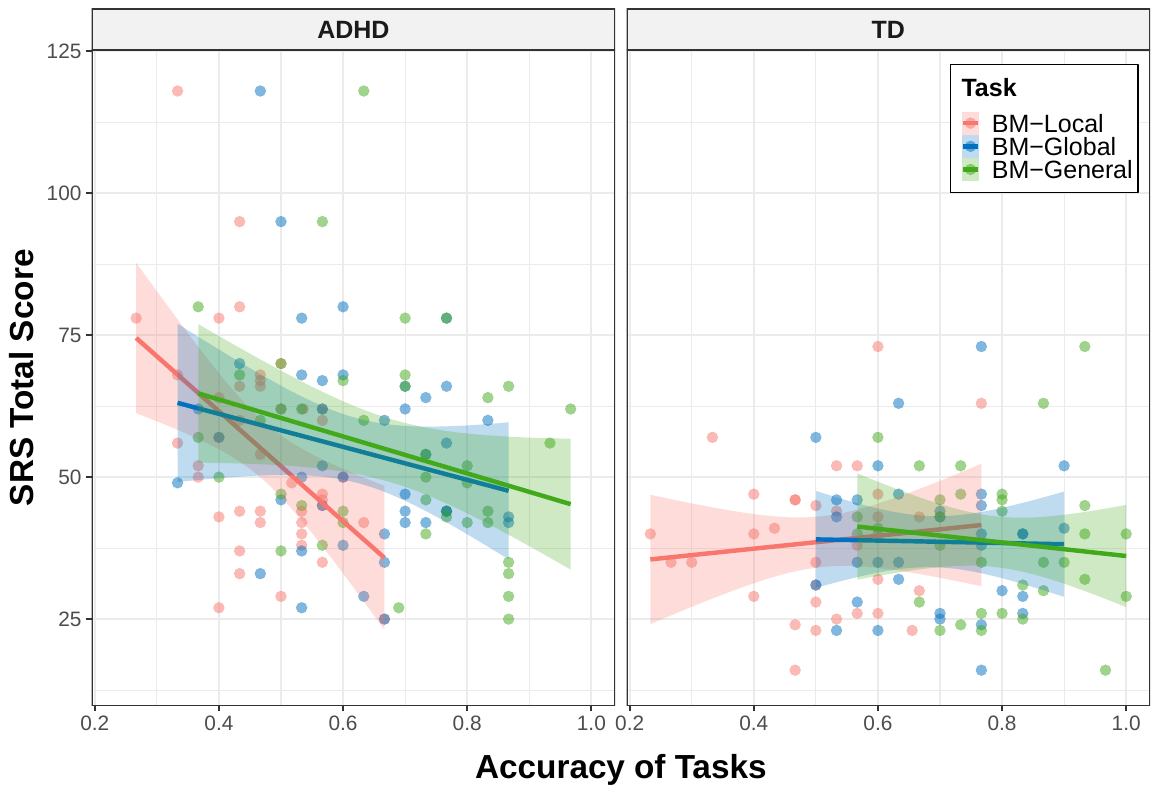

**Figure S2.** **Correlations between the response accuracies and SRS total score.**

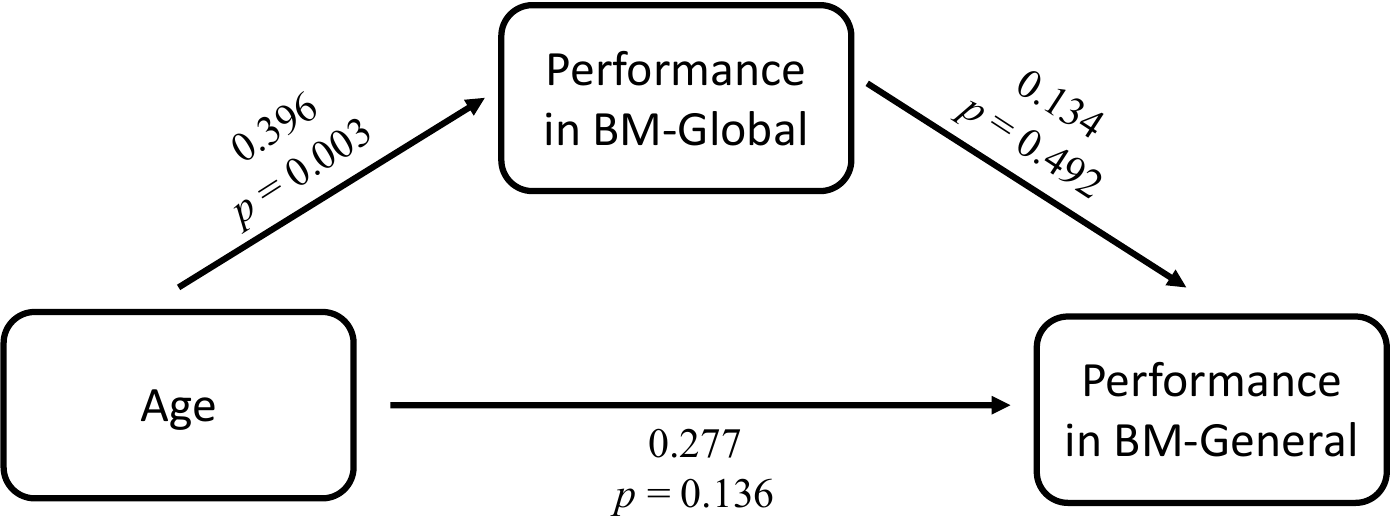

**Figure S3. Factors influencing BM perception in TD children.**

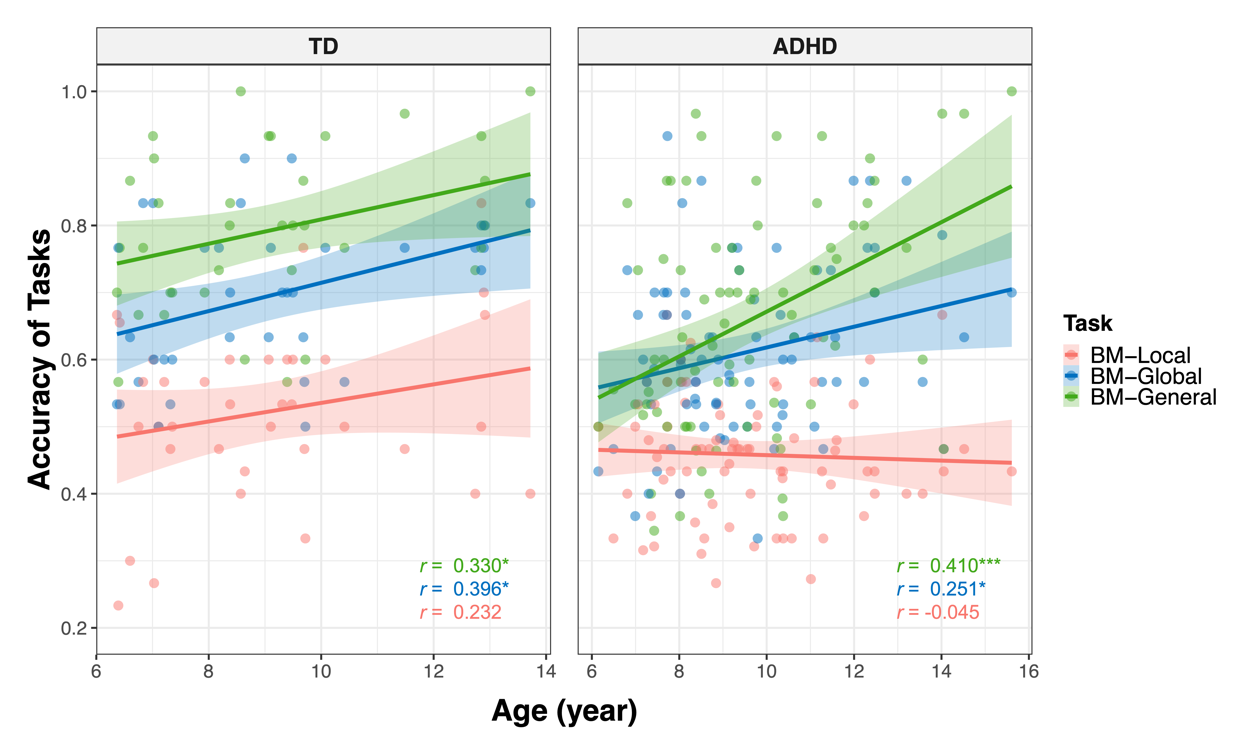

**Figure S4.** **Correlations between the response** **accuracies and age** **(showed by group).** * non-corrected *p* < 0.01, *** non-corrected *p* < 0.001.

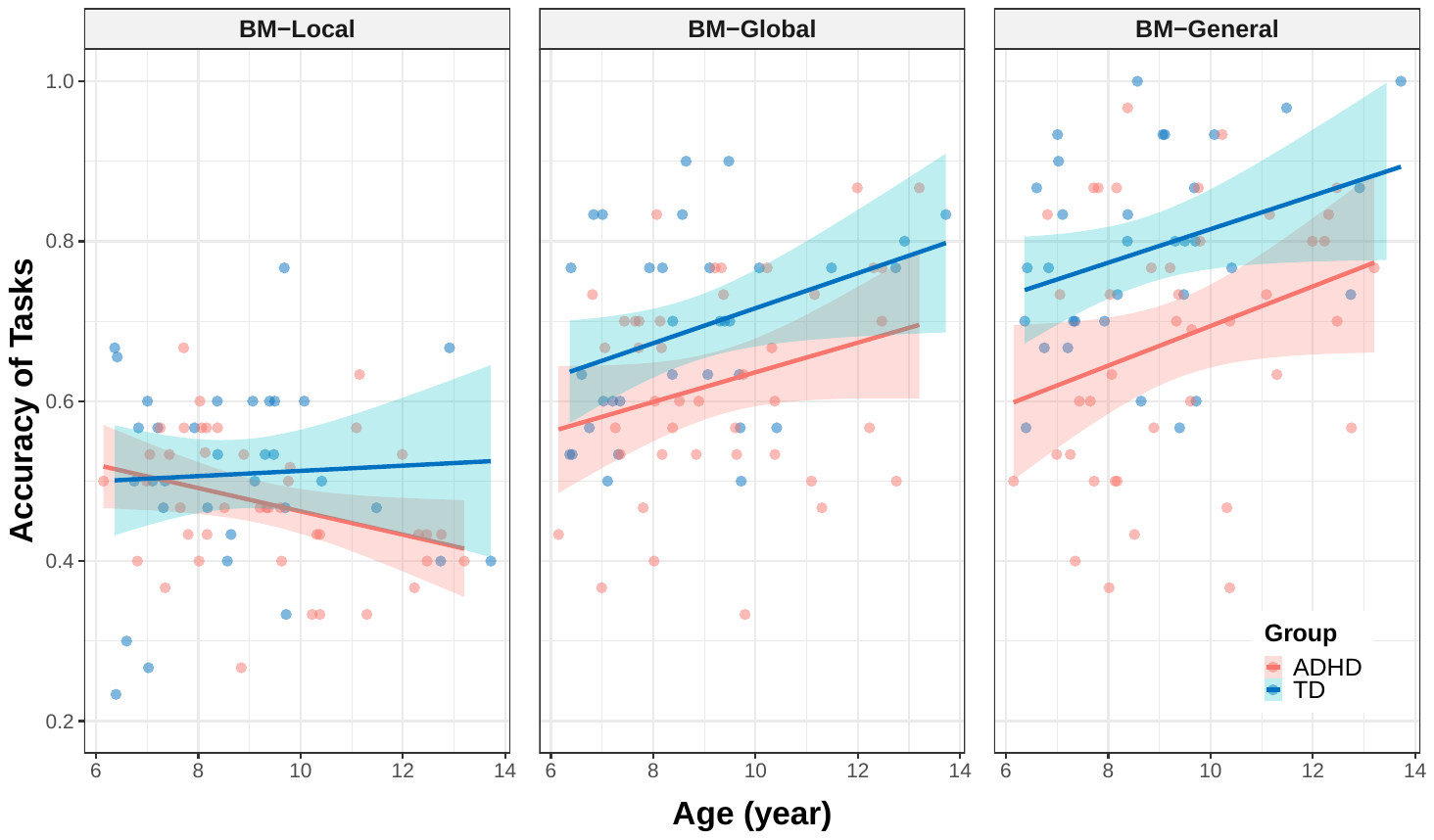

**Figure S5.** **Correlations between the response** **accuracies and age (showed by task).** * non-corrected *p* < 0.01, *** non-corrected *p* < 0.001.

**Tables**

**Table S1. Coefficients and summaries of models a-f**

| Model | Predictors | Standardized Coefficients | 95 % Confidence Interval | T Statistics | P Value | R Square | Std. Error of the Estimate^a^ |
| --- | --- | --- | --- | --- | --- | --- | --- |
| Model a | No significant variable | — | — | — | — | — | — |
| Model b | *age* | 0.251 | [0.033, 0.469] | 2.289 | 0.025 | 0.063 | 0.974 |
| Model c | *age* | 0.365 | [0.172, 0.559] | 3.759 | < 0.001 | 0.339 | 0.829 |
|  | *Perceptual Reasoning* | 0.271 | [0.082, 0.459] | 2.862 | 0.005 |  |  |
|  | *ACC_BM-Global_* | 0.290 | [0.097, 0.484] | 2.987 | 0.004 |  |  |
| Model e | No significant variable | — | — | — | — | — | — |
| Model f | *age* | 0.396 | [0.076, 0.716] | 2.515 | 0.017 | 0.157 | 0.932 |
| Model g | *age* | 0.330 | [0.001, 0.659] | 2.039 | 0.049 | 0.109 | 0.958 |

**Table S2. Coefficients and summaries of models for the relationship between SRS total score and BM processing**

| Model | Predictors | Standardized Coefficients | 95 % Confidence Interval | T Statistics | P Value | R Square | Std. Error of the Estimate^a^ | |
| --- | --- | --- | --- | --- | --- | --- | --- | --- |
| Model 1^b^: *SRS = β_0_ + β_1_ * ACC_BM-Local_ + β_2_ * D + β_3_ * (ACC_BM-Local_ * D)* | *ACC_BM-Local_* | 0.066 | [-0.190, 0.322] | 0.513 | 0.609 | 0.328 | | 0.837 |
|  | *D* | 0.821 | [0.427, 1.215] | 4.151 | < 0.001 |  |  |  |
|  | *ACC_BM-Local_ * D* | -0.629 | [-1.030, -0.228] | -3.127 | 0.003 |  |  |  |
| Model 2^b^: *SRS = β_0_ + β_1_ * ACC_BM-Global_ + β_2_ * D + β_3_ * (ACC_BM-Global_ * D)* | *ACC_BM-Global_* | -0.015 | [-0.360, 0.329] | -0.090 | 0.929 | 0.226 | | 0.898 |
|  | *D* | 0.845 | [0.413, 1.277] | 3.898 | < 0.001 |  |  |  |
|  | *ACC_BM-Global_ * D* | -0.195 | [-0.636, 0.246] | -0.881 | 0.381 |  |  |  |
| Model 3^b^: *SRS = β_0_ + β_1_ * ACC_BM-General_ + β_2_ * D + β_3_ * (ACC_BM-General_ * D)* | *ACC_BM-General_* | -0.104 | [-0.498, 0.290] | -0.526 | 0.600 | 0.251 | | 0.883 |
|  | *D* | 0.765 | [0.318, 1.212] | 3.411 | 0.001 |  |  |  |
|  | *ACC_BM-General_ * D* | -0.179 | [-0.653, 0.296] | -0.751 | 0.455 |  |  |  |
| Model 4^c^: *SRS_ADHD_ = β_0_ + β_1_ * ACC + β_2_ * D_1_ + β_3_ * D_2_ + β_4_ * (ACC * D_1_) + β_5_ * (ACC * D_2_)* | *ACC* | -0.828 | [-1.357, -0.299] | -3.098 | 0.002 | 0.112 | | 0.961 |
|  | *D_1_* | 0.683 | [0.100, 1.267] | 2.317 | 0.022 |  |  |  |
|  | *D_2_* | 0.785 | [0.185, 1.386] | 2.589 | 0.011 |  |  |  |
|  | *ACC * D_1_* | 0.580 | [-0.057, 1.216] | 1.803 | 0.074 |  |  |  |
|  | *ACC * D_2_* | 0.550 | [-0.052, 1.152] | 1.808 | 0.073 |  |  |  |

a. Std. Error of the Estimate is the standard deviation of the error term, and is the square root of the Mean Square Residual (or Error)

b. The variable *D* was a dummy variable (TD group: *D* = 0, ADHD group: *D* = 1, i.e., TD group as a reference).

c. The variable *D_1_* and *D_2_* were dummy variables (BM-Local: *D_1_* = 0, *D_2_* = 0; BM-Global: *D_1_* = 1, *D_2_* = 0; BM-General: *D_1_* = 0, *D_2_* = 1, i.e., BM-Local as a reference).

**Table S3. Coefficients and summaries of models for the relationship between age and BM processing**

| Model | Predictors | Standardized Coefficients | 95% Confidence Interval | T Statistics | P Value | R Square | Std. Error of the Estimate^a^ | |
| --- | --- | --- | --- | --- | --- | --- | --- | --- |
| Model 5^b^: *ACC_ADHD_ = β_0_ + β_1_ * age + β_2_ * D_1_ + β_3_ * D_2_ + β_4_ * (age * D_1_) + β_5_ * (age * D_2_)* | *age* | -0.027 | [-0.203, 0.150] | -0.296 | 0.768 | 0.373 | | 0.801 |
|  | *D_1_* | 0.977 | [0.728, 1.227] | 7.722 | 0.000 |  |  |  |
|  | *D_2_* | 1.269 | [1.019, 1.518] | 10.023 | 0.000 |  |  |  |
|  | *age * D_1_* | 0.228 | [-0.022, 0.478] | 1.800 | 0.073 |  |  |  |
|  | *age * D_2_* | 0.462 | [0.212, 0.712] | 3.639 | < 0.001 |  |  |  |
| Model 6^b^: *ACC_TD_ = β_0_ + β_1_ * age + β_2_ * D_1_ + β_3_ * D_2_ + β_4_ * (age * D_1_) + β_5_ * (age * D_2_)* | *age* | 0.181 | [-0.055, 0.418] | 1.521 | 0.131 | 0.517 | | 0.712 |
|  | *D_1_* | 1.046 | [0.714, 1.379] | 6.236 | < 0.001 |  |  |  |
|  | *D_2_* | 1.636 | [1.303, 1.969] | 9.751 | < 0.001 |  |  |  |
|  | *age * D_1_* | 0.095 | [-0.240, 0.429] | 0.562 | 0.575 |  |  |  |
|  | *age * D_2_* | 0.056 | [-0.278, 0.391] | 0.333 | 0.739 |  |  |  |
| Model 7^c^: *ACC_BM-Local_ = β_0_ + β_1_ * age + β_2_ * D + β_3_ * (age * D)* | *age* | 0.266 | [-0.042, 0.575] | 1.709 | 0.090 | 0.100 | | 0.961 |
|  | *D* | -0.631 | [-1.016, -0.245] | -3.242 | 0.002 |  |  |  |
|  | *age * D* | -0.306 | [-0.684, 0.073] | -1.600 | 0.112 |  |  |  |
| Model 8^c^: *ACC_BM-Global_ = β_0_ + β_1_ * age + β_2_ * D + β_3_ * (age * D)* | *age* | 0.342 | [0.046, 0.638] | 2.290 | 0.024 | 0.172 | | 0.922 |
|  | *D* | -0.723 | [-1.093, -0.353] | -3.874 | < 0.001 |  |  |  |
|  | *age * D* | -0.091 | [-0.454, 0.272] | -0.496 | 0.621 |  |  |  |
| Model 9^c^: *ACC_BM-General_ = β_0_ + β_1_ * age + β_2_ * D + β_3_ * (age * D)* | *age* | 0.229 | [-0.047, 0.505] | 1.643 | 0.103 | 0.280 | | 0.860 |
|  | *D* | -0.884 | [-1.229, -0.540] | -5.081 | < 0.001 |  |  |  |
|  | *age * D* | 0.192 | [-0.146, 0.531] | 1.126 | 0.263 |  |  |  |

a. Std. Error of the Estimate is the standard deviation of the error term, and is the square root of the Mean Square Residual (or Error)

b. The variable *D_1_* and *D_2_* were dummy variables (BM-Local: *D_1_* = 0, *D_2_* = 0; BM-Global: *D_1_* = 1, *D_2_* = 0; BM-General: *D_1_* = 0, *D_2_* = 1, i.e., BM-Local as a reference).

c. The variable *D* was a dummy variable (TD group: *D* = 0, ADHD group: *D* = 1, i.e., TD group as a reference).
